# The role of genetic diversity and arbuscular mycorrhizal fungal diversity in population recovery of the semi-natural grassland plant species *Succisa pratensis*

**DOI:** 10.1101/2021.08.05.455236

**Authors:** Maarten Van Geel, Tsipe Aavik, Tobias Ceulemans, Sabrina Träger, Joachim Mergeay, Gerrit Peeters, Kasper van Acker, Martin Zobel, Kadri Koorem, Olivier Honnay

## Abstract

**Background:** Ecosystem restoration is as a critical tool to counteract the decline of biodiversity and recover vital ecosystem services. Restoration efforts, however, often fall short of meeting their goals. Although functionally important levels of biodiversity can significantly contribute to the outcome of ecosystem restoration, they are often overlooked. One such important facet of biodiversity is within-species genetic diversity, which is fundamental to population fitness and adaptation to environmental change. Also the diversity of arbuscular mycorrhizal fungi (AMF), obligate root symbionts that regulate nutrient and carbon cycles, potentially plays a vital role in mediating ecosystem restoration outcome. In this study, we investigated the relative contribution of intraspecific population genetic diversity, AMF diversity, and their interaction, to population recovery of *Succisa pratensis*, a key species of nutrient poor semi natural grasslands. We genotyped 180 individuals from 12 populations of *S. pratensis* and characterized AMF composition in their roots, using microsatellite markers and next generation amplicon sequencing, respectively. We also investigated whether the genetic makeup of the host plant species can structure the composition of root-inhabiting AMF communities.

**Results:** Our analysis revealed that population allelic richness was strongly positively correlated to relative population growth, whereas AMF richness and its interaction with population genetic diversity did not significantly contribute. The variation partitioning analysis showed that, after accounting for soil and spatial variables, the plant genetic makeup explained a small but significant part of the unique variation in AMF communities.

**Conclusions:** Our results confirm that population genetic diversity can contribute to population recovery, highlighting the importance of within-species genetic diversity for the success of restoration. We could not find evidence, however, that population recovery benefits from the presence of more diverse AMF communities. Our analysis also showed that the genetic makeup of the host plant structured root-inhabiting AMF communities, suggesting that the plant genetic makeup may be linked to genes that control symbiosis development.

## Background

Terrestrial biodiversity is increasingly threatened by land-use change, climate change, pollution and invasive species (1–3), potentially compromising its contribution to resilient provisioning of ecosystem functions and services, such as biomass production, air and water quality modulation, carbon sequestering, soil formation, pollination and nutrient cycling (4, 5). Ecosystem restoration has now emerged as a critical tool to counteract the decline of biodiversity and to recover vital ecosystem services (6). Recently, the United Nations declared 2021-2030 as the decade of ecosystem restoration to protect and revive ecosystems all around the world and committed to restore over 350 million hectares of degraded land (UN 2019, EU Biodiversity Strategy 2030). Although such initiatives have high potential, restoration efforts to protect and regain biodiversity often fall short (7–9) and outcomes of restoration projects differ widely, ranging from near-total success to complete failure (10). Although different strategies to advance ecological restoration success have been proposed (11), the role of functionally important levels of biodiversity have gained little attention.

One such important but often overlooked facet of biodiversity in a restoration context is within-species genetic diversity (12, 13). Genetic diversity is fundamental to population fitness, adaptation to environmental change, habitat loss and disease, with a low genetic diversity increasing the risk of population extinction (14). Furthermore, the role of intraspecific genetic diversity in ecosystem functioning, resilience and service provisioning has been increasingly demonstrated (13, 15, 16). Restoration projects, however, rarely take into account intraspecific population diversity (17) and restoration practitioners often disregard the importance of genetic aspects within their restoration goals (18). Even in the post-2020 global biodiversity framework, a significant global conservation policy mechanism, intraspecific genetic diversity received little attention (19, 20). Therefore, neglecting intraspecific genetic diversity in restoration jeopardizes restoration success, population recovery and the provision of vital ecosystem services.

It is well known that also belowground biodiversity can mediate restoration success, as it shapes aboveground biodiversity, ecosystem functioning and services (21, 22). Key components of belowground biodiversity are arbuscular mycorrhizal fungi (AMF), which are obligate root symbionts, associating with the majority of terrestrial plant species (23). In return for plant assimilated carbon, these fungi provide nutrients, such as nitrogen and phosphorus, to their host plants (23). Thereby AMF regulate nutrient and carbon cycles and play a vital functional role in natural ecosystems (24). With their large surface area of extraradical hyphae, AMF increase root hydraulic conductivity and water absorption, protecting plants against drought stress (25). AMF also defend their hosts against pathogens and increase the tolerance against biotic stresses (26, 27). Additionally, AMF hyphae affect soil structure and aggregate stability, which indirectly increase water availability for the plant (28, 29). Given these properties, AMF diversity potentially plays a pivotal role in mediating ecosystem restoration outcome and population recovery (30). Indeed, mycorrhizal inoculation has been shown to facilitate establishment of vegetation cover and restoration of diverse plant communities (31, 32). Yet, how AMF diversity and community composition mediate restoration outcomes remains understudied.

As a plant’s unique combination of genes, i.e. its genetic makeup, can shape key physiological and morphological plant traits, it can also be expected to affect the function and composition of microorganisms associated to their roots. The genetic makeup of the host plant can indeed alter the composition of rhizosphere fungal communities (33). Inversely, soil microbes can also specialize on specific genotypes within wild plant populations, affecting seedling performance and plant community dynamics (34). Although there is limited information available on the link between plant genetic makeup and AMF communities, there is evidence that plant breeding, which significantly changes the crop genetic makeup, affects also the host-AMF interaction (35). In wheat, for example, it has been demonstrated that cultivars differ in terms of AMF root colonization intensity, nutrient uptake, growth response and root-inhabiting AMF community composition (36–38). Information is, however, lacking about the role of the genetic makeup of the host plant on the structure of root-inhabiting AMF communities in wild plant populations.

In this study, we investigated the relative contribution of intraspecific population genetic diversity, AMF diversity, and their interaction, to population recovery of *Succisa pratensis*, a characteristic plant species of nutrient-poor semi-natural grasslands. We sampled 180 individuals of *S. pratensis* from 12 populations. Plants were genotyped and AMF communities in their roots were characterized using microsatellite markers and next generation amplicon sequencing, respectively. We hypothesized that both plant genetic diversity, AMF diversity and their interaction contribute to population recovery. Additionally, we hypothesized that the genotype of the host plant drives the AMF community composition in plant roots.

## Methods

### Study species

*Succisa pratensis* Moench is an AMF-dependent diploid long-lived perennial rosette herb with a short vertical rhizome (39). It flowers from July to October, producing 1 to 21 flower heads on up to ten flowering stems of 20 to 80 cm. Each flower head contains 70 to 110 violet four-lobed tube flowers (39). As the plant flowers relatively late, it is an important source of nectar and pollen for many insects late in the season (40). Although *S. pratensis* is self-compatible, insect-pollination by bees, bumblebees and hoverflies promotes the seed viability. The seeds fall close to the mother plant and can germinate immediately, but can be dispersed over long distances by epi- and endozoochory. Seeds only survive for a short time and thus do not form a persistent seed bank. Next to sexual reproduction, new rosettes are sporadically formed clonally at the end of short stolons. Historically, *S. pratensis* occurred widely throughout the temperate zones of Eurasia in nutrient-poor acidic and calcareous grasslands, heathlands, unfertilized hay meadows and calcareous fens (39). Due to land use changes, habitat fragmentation and degradation, however, the distribution of *S. pratensis* in the Netherlands declined by 74% since 1935 (40). The remaining populations are often isolated and/or small in size.

### Study area and sampling

This study was conducted in the Hageland region (Flanders, Belgium), which has a maritime mesothermic climate with significant precipitation in all seasons, with an average annual precipitation and temperature of 792 mm and 11.0°C, respectively. On the basis of known population sizes in 2010 (41), we sampled 12 wild populations in September 2020 along a gradient of population recovery rates in terms of population size, following restoration management. Per population, a pooled topsoil sample (0-10 cm) consisting of ten soil cores was collected with an auger of 2 cm diameter. In each population, 15 individuals were randomly selected for sampling. For each individual, a leaf sample and pooled root sample, mainly consisting of fine roots, was collected. Care was taken to leave a sufficient amount of roots and leaves of the plant unharmed to ensure its survival. Leaf and root samples were stored at −20°C until further analysis and soil samples were stored at 4°C for maximum one week to prevent nitrogen loss. In total, 12 soil, 180 leaf and 180 root samples were collected.

### Soil chemical analyses

For each soil sample, soil pH was quantified using a glass electrode in a 1:10 soil/water mixture. As a measure of plant-available soil inorganic nitrogen, ammonium and nitrate were quantified by shaking 10 g of soil in 200 mL of 1 M potassium chloride solution for one hour. Total soil inorganic nitrogen was calculated as the sum of soil ammonium and soil nitrate. As a measure of plant-available soil phosphorus, Resin P values were quantified by shaking 3 g of soil in 30 mL water for 16 hours with anion exchange membranes and subsequent colorimetric analysis of the extracts using the Malachite Green reaction. Extracts were analyzed colorimetrically using the Evolution 201 UV-visible Spectrophotometer (Thermo Scientific, Waltham, MA, USA). Moisture content was determined by the weight loss of 10 g of fresh soil after evaporation of water content in an oven at 105 °C for 1 day. Organic matter was quantified by the weight loss of 10 g of dry soil after combustion of organic matter at 700 °C.

### Genotyping *Succisa pratensis* individuals

Per leaf sample of *S. pratensis*, DNA was extracted from 50 mg of leaf material with the DNA Isolation Plus Kit (Norgen Biotek, Canada) following the manufacturer’s protocol. The genetic variation of *S. pratensis* was assessed using microsatellite analysis (Single Sequence Repeats, SSR), as they are widely applied for genotyping plants, highly informative and cost-effective (42). Twelve loci developed by the Research Institute for Nature and Forest (Belgium) were amplified (Supr-10, Supr-12, Supr-13, Supr-14, Supr-23, Supr-30, Supr-31, Supr-32, Supr-34, Supr-35, Supr-36 and Supr-43; Supplementary information Table S1) (41). Three multiplex PCRs performed on a Bio-Rad T100 thermal cycler (Bio-Rad Laboratories, CA, USA) were constructed of three to six loci in 10 μl reactions. Each multiplex contained 1 μl template DNA, 50-200 nM primer concentration and 5 μl Qiagen Multiplex PCR Master Mix. The PCR cycling profile consisted of an initial denaturation at 94°C for 2 min, 30 cycles consisting of 45 s at 94°C, 45 s at 57°C and 45 s at 72°C, followed by a final elongation of 10 min at 72°C. Fragments were sized on an ABI Prism 3500 Genetic Analyzer (Applied Biosystems) and scored with Genemapper v6 (Applied Biosystems).

### Characterizing AMF communities using next generation amplicon sequencing

Per root sample, fresh roots with a diameter of 3 mm or less were mixed and DNA was extracted from 100 mg of root material using the UltraClean Plant DNA Isolation Kit (MoBio Laboratories, Solana Beach, CA, USA). All DNA extracts were PCR amplified using the sample-specific barcode-labelled versions of the primers AMV4.5NF and AMDGR (43). This primer pair is AMF specific and consistently characterizes AMF communities based on the most variable part of the small subunit (SSU) rRNA gene region (44). PCR was performed on a Bio-Rad T100 thermal cycler (Bio-Rad Laboratories, CA, USA) in a reaction volume of 20 μl, containing 0.15 mM of each dNTP, 0.5 μM of each primer, 1x Titanium Taq PCR buffer, 1U Titanium Taq DNA polymerase (Clontech Laboratories, Palo Alto, CA, USA), and 1 μl genomic DNA. DNA samples were denatured at 94°C for 2 min. Next, 35 cycles were ran, consisting of 45 s at 94°C, 45 s at 65°C and 45 s at 72°C, followed by a final elongation of 10 min at 72°C. The amplicons were separated by agarose gel electrophoresis, cut out, purified using the Qiaquick gel extraction kit (Qiagen, Hamburg, Germany), and quantified using the Quant-iT PicoGreen dsDNA Assay Kit and Qubit fluorometer (Invitrogen, Ghent, Belgium). Next, they were pooled in equimolar quantities. The final amplicon library was sequenced at the Genomics Core UZ Leuven using an Illumina MiSeq sequencer with v2 500 cycle reagent kit (Illumina, San Diego, CA, USA). Raw Illumina data was uploaded to the Sequence Read Archive (accession number PRJNA725853).

Sequences from the Illumina run, obtained as de-multiplexed FASTQ files, were clustered into operational taxonomic units (OTUs) using USEARCH (v. 11) following the recommended pipeline (45). First, paired-end reads were merged to form consensus sequences using the *fastq_mergepairs* command. Next, quality filtering of the reads was performed with the *fastq_filter* command, allowing a maximum expected error of 0.5 for the individual sequences. Then, sequences were dereplicated and sorted by abundance. Sequences occurring only once in the entire dataset were removed prior to clustering as this has been shown to improve the accuracy of diversity estimates (46). Sequences were clustered into OTUs defined at 97% sequence similarity using the UPARSE algorithm implemented in USEARCH, during which chimeric sequences were also removed (45). OTUs were assigned to a taxonomic identity by querying the representative sequence against the MaarjAM database (47). Taxonomic assignments were considered reliable when ≥ 300 BLAST score was found. Only OTUs identified as Glomeromycota were retained in the dataset.

### Statistics

The relative population growth was calculated as (N_2010_-N_2020_)/N_2010_ where N_2010_ and N_2020_ represent the population size in 2010 and 2020, respectively. To assess the *S. pratensis* intraspecific population genetic diversity, allelic richness and expected heterozygosity were calculated based on the 12 SSR markers. Allelic richness per population was calculated as the average number of alleles across all loci using the *allel.rich* function (R package PopGenReport) (48). Expected heterozygosity per population was calculated as 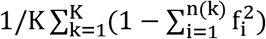, where n(k) is the number of alleles of locus k with a total of K loci and f_i_ is the allele frequency of allele i in a population, using the *Hs* function (R package adegenet) (49).

We calculated AMF richness as the number of AMF OTUs per sample. To prevent bias due to different sequencing depth, resampling techniques to the smallest number of reads per sample are often used. In our AMF dataset, however, the number of reads per sample (median = 23 808) was not related to observed AMF richness (F = 1.07, P = 0.302). Nevertheless, to obtain a reliable and comparable measure of AMF OTU richness and avoid any bias due to sequencing depth, AMF richness was extrapolated or interpolated to 4 000 reads per sample using the *iNEXT* function (R package iNEXT) (50). Population AMF richness was calculated as the average AMF richness of the 15 individuals sampled in the population.

To test the first hypothesis, whether population genetic diversity and AMF diversity positively affect population recovery, a linear model was built using JMP (v15.1) to relate relative population growth to population allelic richness, AMF richness and their interaction. As allelic richness and expected heterozygosity were highly correlated (F = 33.61, P < 0.001), only the former variable was included in the model. The soil variables (moisture content, organic matter, pH, phosphorus and nitrogen content) were also included to account for effects of these variables on the relative population growth. The Akaike Information Criterion (AIC) was used to select the most parsimonious model (i.e. with the lowest AIC) out of a range of reduced models compared with the full model including all explanatory variables, also allowing an interaction effect between population allelic richness and AMF richness. The interaction between population allelic richness and AMF richness was visualized in a contour plot, where the relative growth of the population was represented by the contour curves.

To test the second hypothesis, whether the genotype of a plant can affect AMF community composition, redundancy analysis (RDA) and variation partitioning were performed. First, AMF community composition was directly related to allelic composition using canonical redundancy analysis (RDA) using the *rda* function of the R package vegan. To determine which alleles significantly explained variation in the AMF communities, forward selection was performed using the *ordistep* function (1 000 Monte Carlo permutations, α < 0.05). To test the significance of the selected alleles in the final model, permutation tests on the individual terms (1000 permutations) were performed using the *anova.cca* function. Second, to investigate the relative contribution of plant allelic composition compared to geography and soil chemical variables to explain the total variation of AMF communities, variance partitioning was performed using the *varpart* function of the R package vegan. Three explanatory matrices were used: geography (to account for geographical auto-correlations), soil data (to account for differences in soil chemical variables) and allelic composition. For the geographical matrix, a set of spatial predictors from the geographical coordinates were calculated by principle coordinates of neighbor matrices (PCNM) using the *pcnm* function of the *vegan* package (51, 52). Only the significant explanatory variables, as determined by forward selection *(ordistep* function), in each of the three matrices were included in the variance partitioning. The *venneuler* package in R was used to create Venn diagrams to visually present the results.

## Results

### Relative population growth and genetic diversity

The relative growth of the sampled populations as measured between the year 2010 and 2020 ranged from −0.55 to 8.21 with an average relative growth of 3.09. The microsatellite analysis found a total of 71 alleles across all individuals and the twelve loci. All loci were polymorphic, with the number of alleles ranging from 3 to 13 (mean 5.91) per locus (Supplementary information Table S2). Current population size was not related to allelic richness (F = 1.49, P = 0.251) or expected heterozygosity (F = 1.30, P = 0.281).

### Characterization of AMF communities

Illumina sequencing of the 180 *Succisa pratensis* DNA root samples yielded a total of 4 848 750 AMF sequences and 121 AMF OTUs across seven families (Supplementary information Table S3). The majority of the OTUs belonged to the Glomeraceae (82 OTUs), Claroideoglomeraceae (13 OTUs) and Acaulosporacea (11 OTUs), whereas other AMF families counted less than 10 OTUs. The average AMF richness per population ranged from 9.71 to 34 AMF OTUs (mean 16.11).

### Drivers of population growth

The model selection procedure relating the relative population growth to population allelic richness, AMF richness and their interaction, selected population allelic richness as the only explanatory variable (Fig. 1). The final model had an AIC of 54.02 and an R^2^ adjusted of 0.596 (F = 17.2, P = 0.002), while the full model had an AIC of 275.70. The parameter estimate of the relationship between relative population growth and population allelic richness was 3.75, indicating a positive relationship between both variables. The average population AMF richness and its interaction with population allelic richness were not included in the final model (Fig. 2). Also the soil variables (moisture content, organic matter, pH, phosphorus and nitrogen content) were not included in the final model and thus did not explain variation in relative population growth of *S. pratensis*.

**Figure 1.**
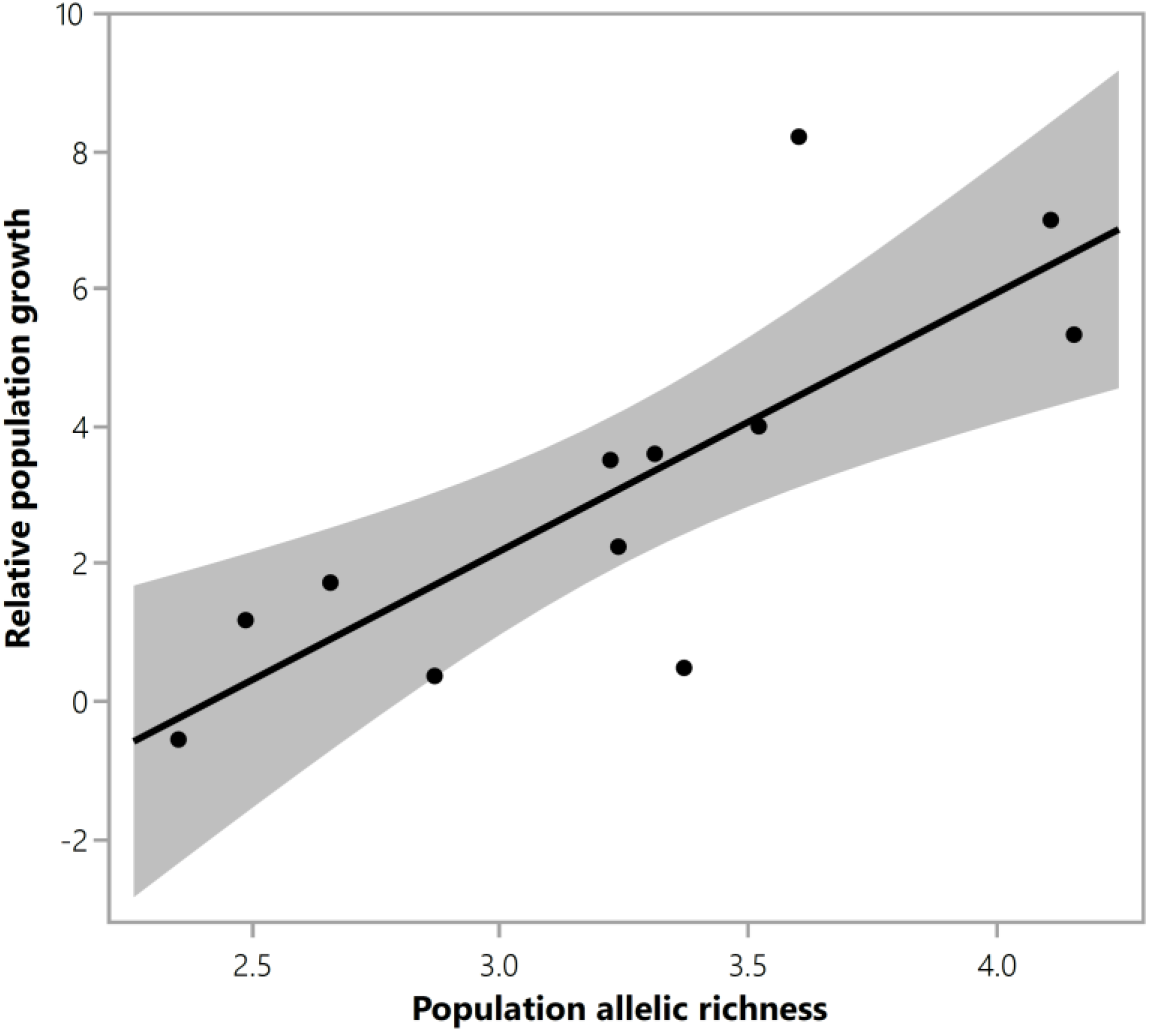
The relationship between relative population growth and population allelic richness of *S. pratensis.* Population allelic richness was the only explanatory variable significantly correlated to relative population growth (R^2^ adjusted of 0.596, F = 17.2, P = 0.002).

**Figure 2.**
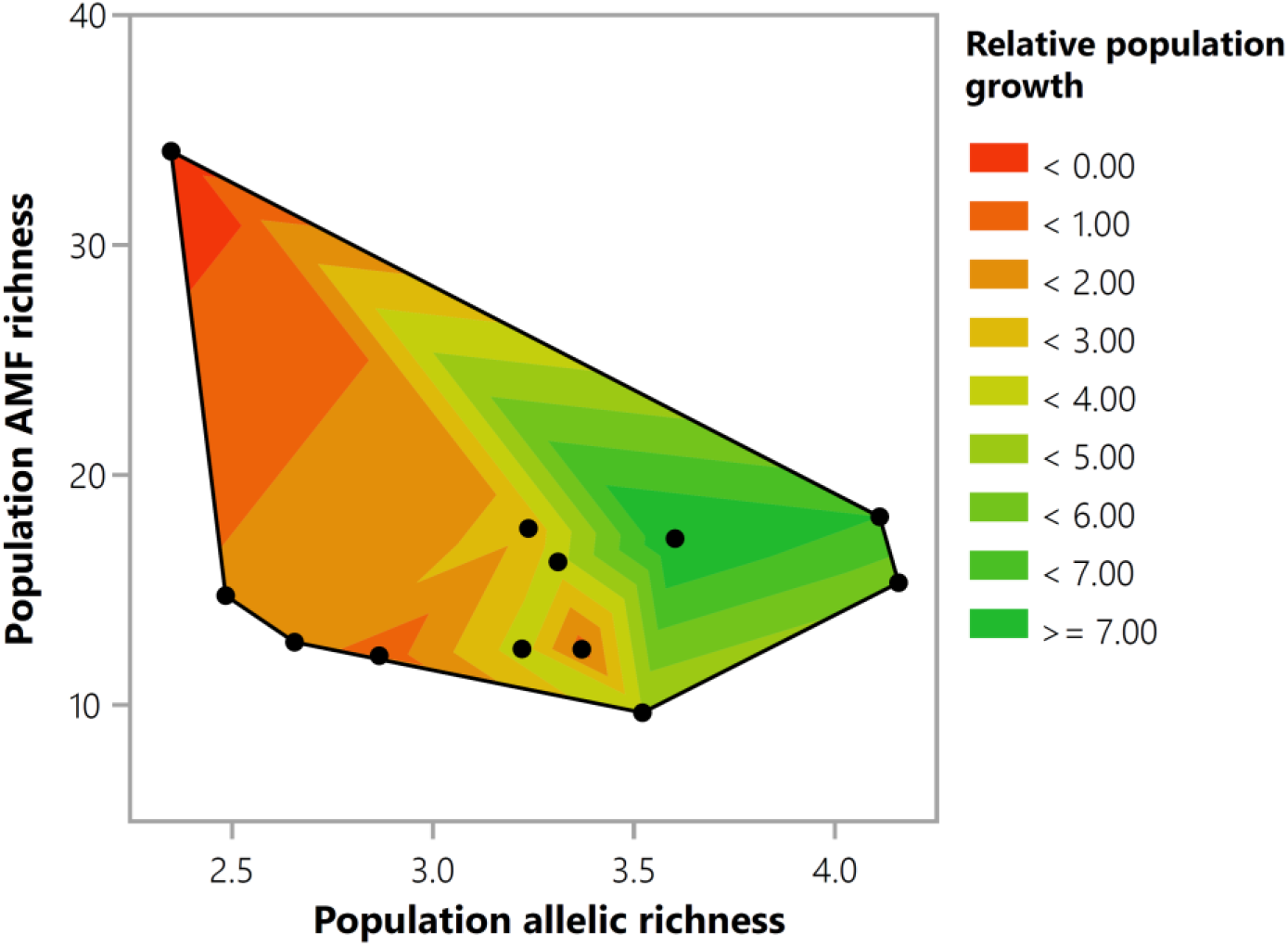
A contour plot visualizing the interaction between population allelic richness and AMF richness of *S. pratensis*. The relative population growth is represented by the contour curves. No significant interaction between population allelic richness and AMF richness was found.

### The relationship between plant genotype and AMF communities

The RDA with forward selection selected 12 alleles significantly explaining 18.1% variation in AMF communities (Table 1). The variation partitioning analysis, to investigate the relative contribution of plant allelic composition compared to geographical and soil chemical variables to explain the total variation of AMF communities, the forward selection procedure selected all four spatial predictor variables of the geographical matrix: PCNM1 (F = 8.07, P < 0.001), PCNM2 (F = 9.91, P < 0.001), PCNM3 (F = 9.21, P < 0.001) and PCNM4 (F = 5.75, P < 0.001). Among the soil data, the forward selection procedure selected all five soil chemical variables significantly explaining variation in AMF communities: moisture content (F = 8.91, P < 0.001), organic matter (F = 10.60, P < 0.001), pH (F = 16.94, P < 0.001), phosphorus (F = 4.92, P < 0.001) and nitrogen content (F = 5.59, P < 0.001). For the allelic composition matrix, the 12 alleles significantly explaining variation in AMF communities were used (Table 1). All three separate explanatory matrices were significantly related to the AMF communities: geography (F = 8.23, P < 0.001), chemical soil variables (F = 9.39, P < 0.001) and allelic composition (F = 4.29, P < 0.001). Comparison of the three different explanatory matrices using variance partitioning revealed that the chemical soil variables and spatial predictors explained most of the unique variation (R^2^ adjusted = 7.61% and 6.63%, respectively), while the allelic composition explained a small but significant part of the unique variation in AMF communities (R^2^ adjusted = 1.95%, F = 1.42, P = 0.005) (Fig. 3). A large part of the variation explained by allelic composition was shared with the chemical soil variables (R^2^ adjusted = 10.23%) and the spatial predictors (R^2^ adjusted = 6.14%).

**Table 1.** Results of the permutation tests of the canonical redundancy analysis (RDA) relating AMF communities to allelic composition of *S. pratensis* (as selected by forward selection). Results are based on 1 000 permutations. The model explained 18.1% of the variation in AMF communities.

| Allele | F | P |
| --- | --- | --- |
| Supr_13.139 | 15.1723 | <0.001 |
| Supr_36.106 | 5.6919 | <0.001 |
| Supr_23.113 | 4.8512 | <0.001 |
| Supr_10.100 | 4.8321 | <0.001 |
| Supr_14.178 | 4.3118 | <0.001 |
| Supr_12.120 | 3.2801 | <0.001 |
| Supr_34.108 | 2.4771 | 0.014 |
| Supr_32.313 | 2.205 | 0.023 |
| Supr_36.107 | 2.1606 | 0.026 |
| Supr_13.135 | 2.1249 | 0.032 |
| Supr_12.124 | 2.1238 | 0.017 |
| Supr_13.133 | 2.2767 | 0.017 |

**Figure 3.**
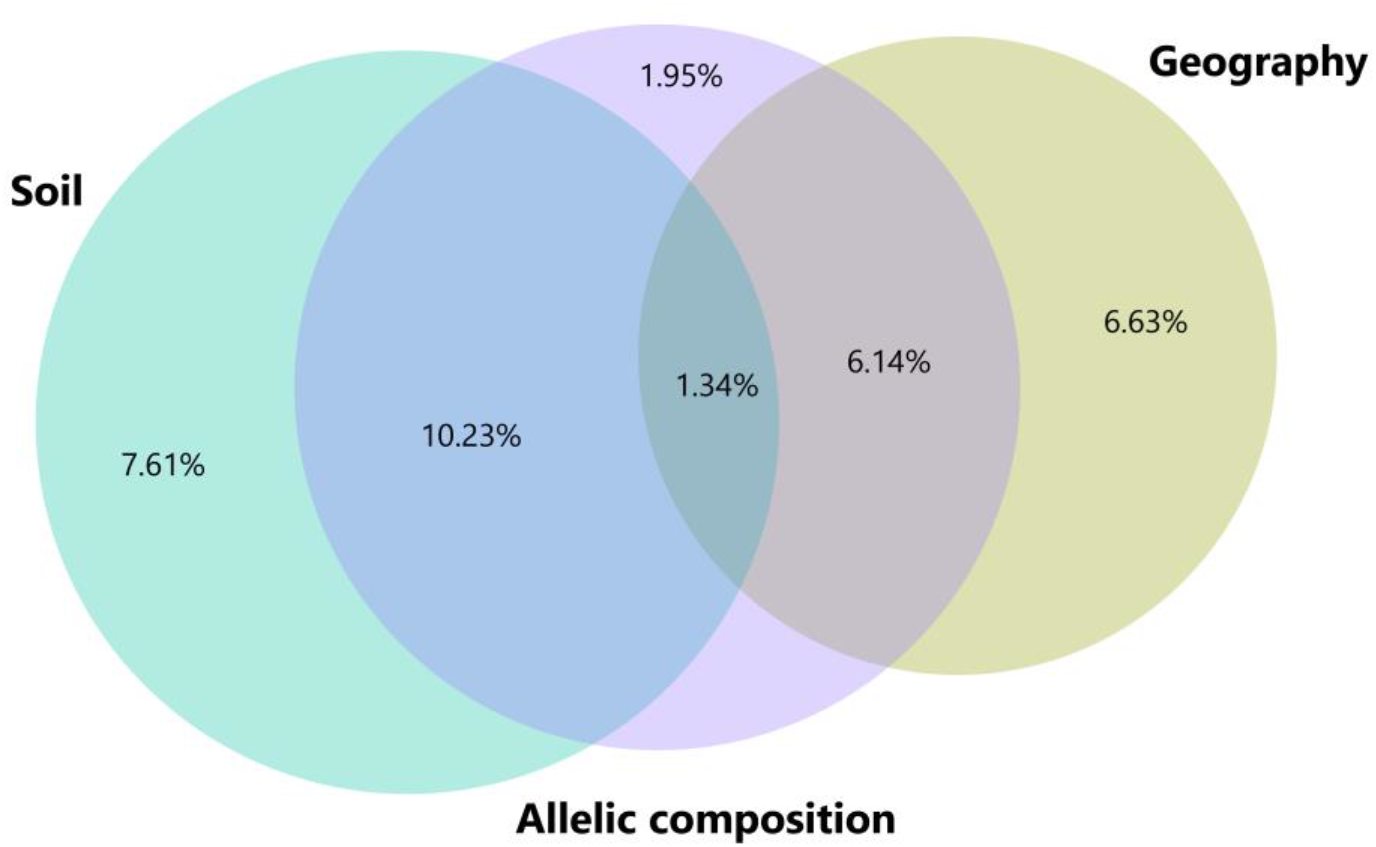
Venn diagram representing variance partitioning results of AMF communities among three explanatory matrices, i.e. geography, soil and allelic composition of *S. pratensis.* The size of the circles is proportional to the variability in AMF communities as explained by a particular explanatory matrix, while overlap of the circles represents the shared variation among explanatory matrices. Numbers indicate the adjusted R^2^ values and thus the variability explained by each partition.

## Discussion

### Plant genetic diversity and population recovery

We tested the hypothesis that plant genetic diversity in combination with AMF diversity contribute to population viability. Our analysis, however, revealed that only population allelic richness was strongly positively correlated to relative population growth, in agreement with the well-known relationship between within-population genetic diversity and population fitness (53). A positive relationship between population genetic diversity and population size was also demonstrated earlier in *S. pratensis* (40). Reduced population genetic diversity can be caused by inbreeding or genetic drift (possibly due to earlier genetic bottlenecks) and diminish the potential to adapt to changing environmental conditions (14). It is also well known that increasing plant population genetic diversity increases seed germination and viability (53, 54), which may explain the observed link between population allelic richness and population recovery of *S. pratensis*.

### AMF diversity and population recovery

Given that AMF play a vital functional role in natural ecosystems as they mediate nutrient and carbon cycles, soil structure and plant interactions with other biota, we expected that AMF diversity would contribute to population recovery (55). Our results, however, showed that both population AMF richness and its interaction with population genetic diversity did not significantly contribute to relative population growth of *S. pratensis.* Therefore, our analysis does not concur with other studies who found that AMF can facilitate restoration of plant populations (31, 32). However, these studies used specific AMF inoculations. The presence of a single specific AMF known to play a functional role for the host plant may thus be more important than diverse AMF communities. In crops, for instance, it has also been shown that specific single species mycorrhizal inoculation is the best approach to enhance plant growth compared to diverse AMF communities (56). Yet, in agreement with our results, another study investigating the effects of AMF on prairie restoration found that AMF did not alter the restoration outcome (57). The absence of a relationship between AMF richness and population recovery may also be explained by the stress that populations which are recovering slowly or are even declining may experience. Stressed host plants allocate more carbon to their root system and symbionts, which may support a more diverse AMF community (58). Indeed, our results showed that the one population that declined also harbored the most diverse AMF communities. This observation is in agreement with Johnson et al. (59) who found that the roots of genetically impoverished plant communities contained more diverse fungal communities compared to genetically rich plant communities. Finally, the sensitivity of next generation amplicon sequencing possibly biased the relationship between AMF diversity and population recovery, as this technique may also detect AMF that are non-functional or even parasitic (60).

### Plant genetic makeup structures associated AMF communities

We hypothesized that the genetic makeup of the host plant can significantly structure root-inhabiting AMF communities in wild plant populations. Indeed, the variation partitioning analysis revealed that, after accounting for soil and spatial variables, the plant genetic makeup explained a small but significant part of the unique variation in AMF communities, thus supporting our hypothesis. The microsatellite markers used to characterize the genetic makeup of the host plant are putatively neutral and hence account for noncoding DNA (61). Yet, our analysis shows a small but significant link between neutral microsatellite markers and the structure of AMF communities. This may be explained by genetic hitchhiking, where the genetic makeup of the host plant is closely linked to other genes affecting the structure of root-inhabiting AMF communities. These genes, for instance, can potentially control symbiosis development, including down-regulation of plant defense reactions, mineral nutrient acquisition and resource trading between partners, allowing the host plant to discriminate the best fungal partners and reward them with carbohydrates (62, 63). Future studies should focus on single-nucleotide polymorphisms that reflect adaptive genetic variation to identify genes that may control the association with root symbionts.

### Implications for ecosystem restoration

Although ecosystem restoration has emerged as a critical tool to counteract the decline of biodiversity and recover vital ecosystem services, restoration efforts often fall short (9). Our results confirm that population allelic richness promotes population recovery, highlighting the importance of within-species genetic diversity for the success of population restoration. Therefore, when reintroducing seeds in the context of ecosystem restoration, our results advocate to mix seeds from a variety of sources to increase the genetic diversity and thus the chance for successful population recovery (64, 65). To safeguard long-term population genetic diversity, restoration efforts also need to take into account the connectivity of populations, as habitat fragmentation has been shown to decrease plant genetic diversity (66). Therefore, our results support ecosystem restoration projects that aim for large and connected populations. Our results, however, do not support that population restoration projects should focus on diverse AMF communities. More important than AMF diversity may be the specific AMF-host plant interaction.

## Conclusions

Our results suggest that population genetic diversity can contribute to population recovery, highlighting the importance of within-species genetic diversity for the success of population restoration. Our study, however, did not reveal positive effects of more diverse AMF communities on the recovery of host plant populations. Our analysis also showed that the genetic makeup of the host plant explained a small but significant part of the unique variation in root-inhabiting AMF communities, suggesting that the plant genetic makeup may be linked to genes that control symbiosis development.

## Declarations

### Ethics approval and consent to participate

Not applicable.

### Consent for publication

Not applicable.

### Availability of data and materials

All data generated or analyzed during this study are included in this published article (Table S2 & S3). Raw Illumina sequence data was uploaded to the Sequence Read Archive (accession number PRJNA725853).

### Authors’ contributions

MVG, MZ and OH designed the research. MVG and TC collected the root and leaf samples. KVA analyzed the soil samples. GP prepared the DNA samples for next generation sequencing. JM performed the genotyping. MVG analyzed the data and wrote the first draft of the manuscript. All authors reviewed and edited the final manuscript. MZ and OH supervised the research.

## Acknowledgements

Not applicable.

## References

1. Pecl GT, Araujo MB, Bell JD, Blanchard J, Bonebrake TC, Chen IC, et al. Biodiversity redistribution under climate change: Impacts on ecosystems and human well-being. Science. 2017;355:1–9.

2. Newbold T, Hudson LN, Arnell AP, Contu S, De Palma A, Ferrier S, et al. Has land use pushed terrestrial biodiversity beyond the planetary boundary? A global assessment. Science. 2016;353:288–91.

3. Diaz S, Settele J, Brondizio ES, Ngo HT, Agard J, Arneth A, et al. Pervasive human-driven decline of life on Earth points to the need for transformative change. Science. 2019;366:1–10.

4. Cardinale BJ, Duffy JE, Gonzalez A, Hooper DU, Perrings C, Venail P, et al. Biodiversity loss and its impact on humanity. Nature. 2012;486:59–67.

5. Hautier Y, Tilman D, Isbell F, Seabloom EW, Borer ET, Reich PB. Anthropogenic environmental changes affect ecosystem stability via biodiversity. Science. 2015;348:336–40.

6. Crouzeilles R, Curran M, Ferreira MS, Lindenmayer DB, Grelle CE, Rey Benayas JM. A global meta-analysis on the ecological drivers of forest restoration success. Nat Commun. 2016;7:1–8.

7. Rey Benayas JM, Newton AC, Diaz A, Bullock JM. Enhancement of biodiversity and ecosystem services by ecological restoration: a meta-analysis. Science. 2009;325:1121–4.

8. Menz MH, Dixon KW, Hobbs RJ. Hurdles and opportunities for landscape-scale restoration. Science. 2013;339:526–7.

9. Stokstad E. Global efforts to protect biodiversity fall short. Science. 2020;369:1418.

10. Suding KN. Toward an era of restoration in ecology: successes, failures, and opportunities ahead. Annual Review of Ecology, Evolution, and Systematics. 2011;42:465–87.

11. Aronson J, Goodwin N, Orlando L, Eisenberg C, Cross AT. A world of possibilities: six restoration strategies to support the United Nation’s Decade on Ecosystem Restoration. Restor Ecol. 2020;28:730–6.

12. Des Roches S, Post DM, Turley NE, Bailey JK, Hendry AP, Kinnison MT, et al. The ecological importance of intraspecific variation. Nat Ecol Evol. 2018;2:57–64.

13. Raffard A, Santoul F, Cucherousset J, Blanchet S. The community and ecosystem consequences of intraspecific diversity: a meta-analysis. Biological Reviews. 2019;94:648–61.

14. Leimu R, Mutikainen P, Koricheva J, Fischer M. How general are positive relationships between plant population size, fitness and genetic variation? J Ecol. 2006;94:942–52.

15. Prieto I, Violle C, Barre P, Durand JL, Ghesquiere M, Litrico I. Complementary effects of species and genetic diversity on productivity and stability of sown grasslands. Nature plants. 2015;1:15033.

16. Oliver TH, Heard MS, Isaac NJB, Roy DB, Procter D, Eigenbrod F, et al. Biodiversity and resilience of ecosystem functions. Trends Ecol Evol. 2015;30:673–84.

17. Harzé M, Monty A, Boisson S, Pitz C, Hermann J-M, Kollmann J, et al. Towards a population approach for evaluating grassland restoration - a systematic review. Restor Ecol. 2018;26:227–34.

18. Mijangos JL, Pacioni C, Spencer PB, Craig MD. Contribution of genetics to ecological restoration. Mol Ecol. 2015;24:22–37.

19. Laikre L, Hoban S, Bruford MW, Segelbacher G, Allendorf FW, Gajardo G, et al. Post-2020 goals overlook genetic diversity. Science. 2020;367:1083–5.

20. Hoban S, Bruford M, D’Urban Jackson J, Lopes-Fernandes M, Heuertz M, Hohenlohe PA, et al. Genetic diversity targets and indicators in the CBD post-2020 Global Biodiversity Framework must be improved. Biol Conserv. 2020;208:1–11.

21. Bardgett RD, van der Putten WH. Belowground biodiversity and ecosystem functioning. Nature. 2014;515:505–11.

22. Bahram M, Hildebrand F, Forslund SK, Anderson JL, Soudzilovskaia NA, Bodegom PM, et al. Structure and function of the global topsoil microbiome. Nature. 2018;560:233–7.

23. Smith SE, Read DJ. Mycorrhizal symbiosis. 3rd ed. Cambridge, UK: Academic Press; 2008.

24. Powell JR, Rillig MC. Biodiversity of arbuscular mycorrhizal fungi and ecosystem function. New Phytol. 2018;220:1059–75.

25. Lehto T, Zwiazek JJ. Ectomycorrhizas and water relations of trees: a review. Mycorrhiza. 2011;21:71–90.

26. Veresoglou S, Rillig M. Suppression of fungal and nematode plant pathogens through arbuscular mycorrhizal fungi. Biol Lett. 2012;8:214–7.

27. Harrier LA, Watson CA. The potential role of arbuscular mycorrhizal (AM) fungi in the bioprotection of plants against soil-borne pathogens in organic and/or other sustainable farming systems. Pest Manage Sci. 2004;60:149–57.

28. Rillig M, Mummey D. Mycorrhizas and soil structure. New Phytol. 2006;171:41–53.

29. van der Heijden MG, Streitwolf-Engel R, Riedl R, Siegrist S, Neudecker A, Ineichen K, et al. The mycorrhizal contribution to plant productivity, plant nutrition and soil structure in experimental grassland. New Phytol. 2006;172:739–52.

30. Asmelash F, Bekele T, Birhane E. The Potential Role of Arbuscular Mycorrhizal Fungi in the Restoration of Degraded Lands. Frontiers in microbiology. 2016;7:1095.

31. Neuenkamp L, Prober SM, Price JN, Zobel M, Standish RJ. Benefits of mycorrhizal inoculation to ecological restoration depend on plant functional type, restoration context and time. Fungal Ecol. 2019;40:140–9.

32. Torrez V, Ceulemans T, Mergeay J, de Meester L, Honnay O. Effects of adding an arbuscular mycorrhizal fungi inoculum and of distance to donor sites on plant species recolonization following topsoil removal. Applied Vegetation Science. 2016;19:7–19.

33. Fernández NV, Marchelli P, Tenreiro R, Chaves S, Fontenla SB. Are the rhizosphere fungal communities of Nothofagus alpina established in two different environments influenced by plant genetic diversity? For Ecol Manage. 2020;473:118269.

34. Eck JL, Stump SM, Delavaux CS, Mangan SA, Comita LS. Evidence of within-species specialization by soil microbes and the implications for plant community diversity. PNAS. 2019;116:7371–6.

35. Martín-Robles N, García-Palacios P, Rodríguez M, Rico D, Vigo R, Sánchez-Moreno S, et al. Crops and their wild progenitors recruit beneficial and detrimental soil biota in opposing ways. Plant Soil. 2020;456:159–73.

36. García de Leon D, Vahter T, Zobel M, Koppel M, Edesi L, Davison J, et al. Different wheat cultivars exhibit variable responses to inoculation with arbuscular mycorrhizal fungi from organic and conventional farms. PLoS One. 2020;15:e0233878.

37. Elliott AJ, Daniell TJ, Cameron DD, Field KJ. A commercial arbuscular mycorrhizal inoculum increases root colonization across wheat cultivars but does not increase assimilation of mycorrhiza-acquired nutrients. Plants, Peope, Planet. 2020;0:1–12.

38. Thirkell TJ, Pastok D, Field KJ. Carbon for nutrient exchange between arbuscular mycorrhizal fungi and wheat varies according to cultivar and changes in atmospheric carbon dioxide concentration. Glob Chang Biol. 2020;26:1725–38.

39. Adams AW. Succisa Pratensis Moench. J Ecol. 1955;43:709–18.

40. Vergeer P, Rengelink R, Copal A, Ouborg NJ. The interacting effects of genetic variation, habitat quality and population size on preformance of *Succisa pratensis*. J Ecol. 2003;91:18–26.

41. Baumers M. Genetische effecten van habitatfragmentatie in een achteruitgaande graslandplant: Blauwe knoop (Succisa pratensis). KU Leuven Masterthesis. 2011.

42. Vieira MLC, Santini L, Diniz AL, Munhoz CD. Microsatellite markers: what they mean and why they are so useful. Genet Mol Biol. 2016;39:312–28.

43. Sato K, Suyama Y, Saito M, Sugawara K. A new primer for discrimination of arbuscular mycorrhizal fungi with polymerase chain reaction-denature gradient gel electrophoresis. Grassland Science. 2005;51:179–81.

44. Van Geel M, Busschaert P, Honnay O, Lievens B. Evaluation of six primer pairs targeting the nuclear rRNA operon for characterization of arbuscular mycorrhizal fungal (AMF) communities using 454 pyrosequencing. J Microbiol Methods. 2014;106:93–100.

45. Edgar R. UPARSE: highly accurate OTU sequences from microbial amplicon reads. Nat Methods. 2013;10:996–8.

46. Brown SP, Veach AM, Rigdon-Huss AR, Grond K, Lickteig SK, Lothamer K, et al. Scraping the bottom of the barrel: are rare high throughput sequences artifacts? Fungal Ecol. 2015;13:221–5.

47. Öpik M, Vanatoa A, Vanatoa E, Moora M, Davison J, Kalwij J, et al. The online database MaarjAM reveals global and ecosystemic distribution patterns in arbuscular mycorrhizal fungi (Glomeromycota). New Phytol. 2010;188:223–41.

48. Adamack AT, Gruber B. PopGenReport: simplifying basic population genetic analyses in R. Methods in Ecology and Evolution. 2014;5:384–7.

49. Jombart T, Ahmed I. adegenet 1.3-1: new tools for the analysis of genome-wide SNP data. Bioinformatics. 2011;27:3070–1.

50. Hsieh TC, Ma KH, Chao A. iNEXT: an R package for rarefaction and extrapolation of species diversity (Hill numbers). Methods in Ecology and Evolution. 2016;7:1451–6.

51. Borcard D, Legendre P. All-scale spatial analysis of ecological data by means of principal coordinates of neighbour matrices. Ecol Model. 2002;153:51–68.

52. Borcard D, Legendre P, Avois-Jacquet C, Tuomisto H. Dissecting the spatial structure of ecological data at multiple scales. Ecology. 2004;85:1826–32.

53. Reed DH, Frankham R. Correlation between fitness and genetic diversity. Conserv Biol. 2003;17:230–7.

54. Booy G, Hendriks RJJ, Smulders MJM, Van Groenendael JM, Vosman B. Genetic diversity and the survival of populations. Plant Biol. 2000;2:379–95.

55. Tedersoo L, Bahram M, Zobel M. How mycorrhizal associations drive plant population and community biology. Science. 2020;367:eaba1223.

56. Van Geel M, De Beenhouwer M, Lievens B, Honnay O. Crop-specific and single-species mycorrhizal inoculation is the best approach to improve crop growth in controlled environments. Agronomy for Sustainable Development. 2016;36:37–47.

57. White JA, Tallaksen J, Charvat I. The effects of arbuscular mycorrhizal fungal inoculation at a roadside prairie restoration site. Mycologia. 2008;100:6–11.

58. Werner GD, Kiers ET. Partner selection in the mycorrhizal mutualism. New Phytol. 2015;205:1437–42.

59. Johnson D, Anderson IC, Williams A, Whitlock R, Grime JP. Plant genotypic diversity does not beget root-fungal species diversity. Plant Soil. 2010;336:107–11.

60. Johnson NC, Wilson GWT, Wilson JA, Miller RM, Bowker MA. Mycorrhizal phenotypes and the law of the minimum. New Phytol. 2015;205:1473–84.

61. Holderegger R, Kamm U, Gugerli F. Adaptive vs. neutral genetic diversity: implications for landscape genetics. Landscape Ecol. 2006;21:797–807.

62. Kiers ET, Duhamel M, Beesetty Y, Mensah JA, Franken O, Verbruggen E, et al. Reciprocal rewards stabilize cooperation in the mycorrhizal symbiosis. Science. 2011;333:880–2.

63. Johnson D, Martin F, Cairney JWG, Anderson IC. The importance of individuals: intraspecific diversity of mycorrhizal plants and fungi in ecosystems. New Phytol. 2012;194:614–28.

64. Wilkinson DM. Is local provenance important in habitat creation? J Appl Ecol. 2001;38:1371–3.

65. Bucharova A, Bossdorf O, Hölzel N, Kollmann J, Prasse R, Durka W. Mix and match: regional admixture provenancing strikes a balance among different seed-sourcing strategies for ecological restoration. Conserv Genet. 2019;20:7–17.

66. González AV, Gómez-Silva V, Ramírez MJ, Fontúrbel FE. Meta-analysis of the differential effects of habitat fragmentation and degradation on plant genetic diversity. Conserv Biol. 2020;34:711–20.

